# TracePheno Enables Function-First Inference of Trace-Element Phenotypes from Microbiome Profiles

**DOI:** 10.64898/2026.03.15.711888

**Authors:** Jian Zhou

**Affiliations:** International Institutes of Medicine, Zhejiang University, Yiwu, China

**Keywords:** microbiome, phenotype analysis, trace elements, PICRUSt2, KEGG Orthology, trait inference

## Abstract

Microbiome phenotype analysis usually captures broad organism-level traits, yet clinically and ecologically important programmes for trace-element acquisition, storage, detoxification, and cofactor biosynthesis remain difficult to summarize because the underlying loci are often strain-variable and only partly reflected by taxonomy. We present TracePheno, a function-first framework for inferring microbial phenotypes related to eight common trace elements from gene- or KO-level evidence. The current phenotype panels span iron, zinc, manganese, copper, cobalt/vitamin B_12_, nickel, molybdenum, and selenium. The implementation combines curated core/accessory/ambiguous marker tiers, cohort-invariant bounded support transforms, deterministic core-gated calling, presence/absence-oriented genome trait inference, and a publication-oriented visualization bundle. The bundled release covers ten phenotype panels and three complementary workflows: direct scoring of function matrices, genome-to-trait construction, and taxon-abundance scoring with a precomputed trait matrix. Using the current release, we analysed two local demonstrations that were regenerated for this manuscript. In 11 representative human-gut genomes from the MGnify catalogue, copper homeostasis/resistance and iron acquisition were the most prevalent high-scoring programmes, whereas Firmicutes in this small panel showed stronger cobalamin biosynthesis and selenium-utilization signals than Bacteroidota. In a PICRUSt2-style KO example, zinc acquisition was higher in the case group, whereas iron acquisition, corrinoid transport/cobalt uptake, and selenium utilization were relatively higher in controls. Together, these analyses show that TracePheno can convert genome annotations and predicted KO tables into interpretable, publication-ready trace-element phenotype landscapes while keeping the decision rules explicit, portable, and biologically constrained.

## 1 Introduction

Microbiome workflows now routinely generate taxonomic profiles, gene-family abundance tables, genome annotations, and pathway predictions. Translating those outputs into interpretable phenotypes remains difficult, however, particularly when the phenotypes of interest are mechanistically specific rather than broad descriptors of cell structure or lifestyle. Existing microbial trait resources and phenotype-prediction tools have been most successful for organism-level characteristics such as aerobicity, Gram status, motility, or biofilm formation [Weimann et al., 2016, Madin et al., 2020]. Trace-element metabolism is a harder target because transporters, detoxification systems, siderophore modules, corrinoid pathways, and selenium-related functions can vary strongly below the species level and are often shaped by horizontal gene transfer.

This problem matters in host-associated and environmental microbiology alike. Iron, zinc, manganese, copper, nickel, molybdenum, and selenium participate in nutritional immunity, oxidative stress defence, respiration, metalloenzyme maturation, and corrinoid-dependent metabolism [Hood and Skaar, 2012, Argüello et al., 2013, Juttukonda and Skaar, 2015, Degnan et al., 2014]. A phenotype framework focused on these elements should therefore be anchored to functional evidence rather than to taxonomy alone. KEGG provides pathway and KO structure [Kanehisa et al., 2022]; BacMet captures the broader logic of curated metal-related gene inventories [Pal et al., 2013]; and genome-scale trait tools such as Traitar demonstrate how genome features can be translated into phenotype calls [Weimann et al., 2016]. What has been missing is a dedicated analysis layer that turns these ingredients into a trace-element phenotype workflow for microbiome datasets.

The need is amplified by heterogeneous data types. Shotgun metagenomes and genome annotations can provide direct KO or gene-family evidence, whereas many cohort studies still begin with 16S amplicon data and therefore rely on function prediction. PICRUSt2 remains a widely used route from marker-gene surveys to predicted KEGG Orthology abundance tables [Douglas et al., 2020]. A practical phenotype tool should therefore support both direct function matrices and PICRUSt2-derived KO predictions while keeping the phenotype model itself stable.

Here we present TracePheno, a function-first framework for trace-element phenotype inference. The current release combines a curated phenotype library, deterministic core-gated scoring, optional quality-aware handling of incomplete genomes, and a publication-oriented visualization bundle. We illustrate the framework using two demonstrations generated with the local release: 11 representative human-gut genomes linked to a recent MGnify-based metagenome-assembled genome study [Almeida et al., 2020, Wang et al., 2025], and a compact PICRUSt2-style KO matrix used to emulate a 16S-derived entry point.

## 2 Materials and Methods

### 2.1 Phenotype library design

The bundled TracePheno library currently covers ten phenotype panels: iron acquisition, iron storage/homeostasis, zinc acquisition, manganese acquisition, copper homeostasis/resistance, cobalamin biosynthesis, cobalamin transport/cobalt uptake, nickel utilization, molybdenum cofactor biosynthesis, and selenium utilization. Marker sets were curated from KEGG module logic, KO relationships, the BacMet concept of metal-related resistance curation, and element-specific microbiology reviews that emphasize nutritional immunity, metal transport, cofactor biosynthesis, and homeostasis [Pal et al., 2013, Hood and Skaar, 2012, Argüello et al., 2013, Juttukonda and Skaar, 2015, Degnan et al., 2014, Kanehisa et al., 2022]. Markers were partitioned into three evidence tiers: *core, accessory*, and *ambiguous*. Core markers represent phenotype-defining functions, accessory markers broaden coverage without replacing core evidence, and ambiguous markers are retained at reduced influence when they are biologically relevant but not specific enough to support a call on their own.

### 2.2 Scoring strategy

TracePheno operates on a feature-by-sample matrix in which rows are gene names, KO identifiers, or mixed annotation strings and columns are samples, genomes, or taxa depending on the workflow. Each phenotype consists of weighted marker sets, and each marker set can require either all members or a minimum number of hits. Feature matching is alias-aware and accepts explicit gene symbols, KO identifiers, and mixed annotation strings.

For each marker set *m* in sample *s*, the algorithm first computes coverage *C*_*m,s*_ from the set logic. Matched abundance is then transformed by a bounded square-root term so that support remains sample-intrinsic rather than cohort-relative:

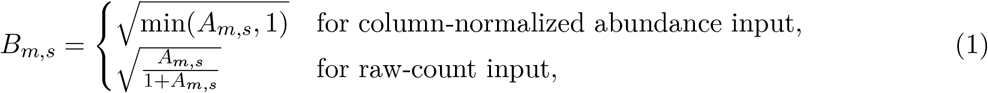

where *A*_*m,s*_ is the summed abundance of matched rows. The bounded support term is

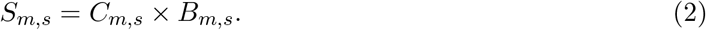

For abundance-aware scoring, marker-set evidence is then

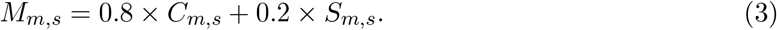

For genome trait inference, TracePheno instead uses binary feature mode, so marker-set evidence reduces to the observed coverage term.

Phenotype scores are not derived from data-dependent threshold scans. Instead, the current release uses deterministic core-gated aggregation. If a phenotype contains core marker sets, the phenotype score is

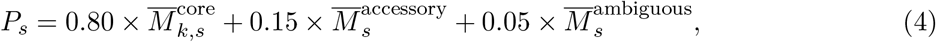

where 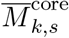 is the mean of the top *k* core marker-set scores and *k* is the phenotype-specific parameter `core_min_hits`. Binary calls are assigned only when the phenotype score exceeds the phenotype-specific threshold hint and at least *k* core marker sets are fully satisfied. This rule is intentionally stricter for multi-block pathways such as cobalamin biosynthesis (`core_min_hits=2`) and molybdenum cofactor biosynthesis (`core_min_hits=3`). If a phenotype lacks core tiers, a deterministic threshold-only rule is used.

### 2.3 Genome quality-aware trait inference

Genome-to-trait inference uses presence/absence-oriented evidence rather than rewarding copy number. When completeness and contamination metadata are available, TracePheno can interpret genome quality using conservative MIMAG-inspired cutoffs [Bowers et al., 2017]. Genomes with ≥ 90% completeness and ≤ 5% contamination are treated as high quality; genomes with ≥ 50% completeness and ≤ 10% contamination are treated as medium quality; and the remainder are classified as low quality. In this framework, negative calls from medium-quality genomes can be flagged as uncertain absences, and calls from low-quality genomes can be withheld rather than forced to zero. This design is meant to prevent missing evidence in incomplete genomes from being overinterpreted as true phenotype absence.

### 2.4 Supported workflows

TracePheno currently supports three major analysis routes.

1. **Function-matrix scoring**: a KO/gene abundance matrix is scored directly to produce phenotype scores, coverage, marker contributions, and group statistics.
2. **Genome-to-trait construction**: a genome annotation matrix is converted into a trait matrix so that downstream phenotype abundance can be estimated from taxon abundance profiles.
3. **Taxon scoring**: a taxon abundance matrix is multiplied by a trait matrix to produce community-level phenotype abundance through trait-weighted aggregation.

If trait matrices contain missing values, the aggregation step can track lower bounds, upper bounds, and knowledge coverage instead of silently coercing missing traits to zero. In addition, the current implementation provides an explicit PICRUSt2 route that reads `pred_metagenome_ unstrat.tsv.gz` or `pred_metagenome_contrib.tsv.gz` style KO tables. The unstratified table is treated as a KO-by-sample matrix, whereas contribution tables are automatically collapsed to sample-level KO abundance. This design lets TracePheno accept 16S-derived KO predictions without changing the phenotype model itself.

### 2.5 Visualization and publication bundle

Beyond conventional heat maps, TracePheno generates a publication-oriented visualization layer including a phenotype landscape plot, clustered phenotype atlas, raincloud-style distribution panels, group trajectory plots, sample embeddings, evidence-tier summaries, marker-contribution panels, and a differential phenotype map for pairwise comparison. The current release packages these plots into composite multi-panel figures and draft legends using a restrained, manuscript-oriented style intended to resemble a high-end journal figure workflow rather than an exploratory dashboard. The figures included in this manuscript were regenerated directly from the current local release immediately before compilation so that the narrative text, numerical summaries, and visual outputs remain synchronized.

### 2.6 Demonstration datasets

We used two demonstrations generated with the current local release. The first was a genome-level run based on the MGnify human-gut v2.0.2 species-representative catalogue, a large reference resource for human gut genomes [Almeida et al., 2020]. For a reproducible case study, we selected 11 representative genomes from the data resources linked by Wang et al. in their humangut metagenome-assembled genome analysis [Wang et al., 2025]. eggNOG annotation tables were parsed into a 7,731-feature by 11-genome matrix. Genome completeness ranged from 97.9% to 100%, and contamination ranged from 0% to 0.41%. The second demonstration was a compact PICRUSt2-style KO matrix with four samples split into case and control groups, used to exercise the 16S-compatible route. Both result directories, summary tables, and publication figures cited below were regenerated from the same code state used to compile this manuscript.

## 3 Results

### 3.1 Deterministic core gating defines the current phenotype model

The current TracePheno release no longer derives binary calls from cohort-dependent threshold scans. Instead, all ten bundled phenotypes are now called with deterministic core-gated rules. The exported threshold file for both demonstrations showed a fixed rule for every phenotype, with phenotype-specific requirements for the number of core blocks that must be satisfied. The strictest bundled examples were cobalamin biosynthesis, which required two core marker-set blocks, and molybdenum cofactor biosynthesis, which required three. This design made the phenotype calls more interpretable because the decision boundary remained attached to pathway logic rather than to the composition of the current dataset.

### 3.2 TracePheno emphasizes function-grounded trace-element phenotypes

The conceptual consequence of this design is that TracePheno defaults to function-grounded inference for trace-element traits rather than transferring phenotypes from taxonomy alone. This distinction is especially relevant for siderophore systems, metal export modules, corrinoid transport, and selenium-related functions, all of which can vary substantially among closely related strains. The current release exposes both continuous scores and binary calls, as well as per-marker evidence and tier-aware summaries.

Table 1 summarizes the phenotype panels currently available in the bundled release.

**Table 1:** Phenotype panels currently implemented in TracePheno.

| Phenotype | Element | Representative marker logic |
| --- | --- | --- |
| Iron acquisition | Iron | Ferrous uptake, ferric-siderophore transport, siderophore biosynthesis, and heme-linked scavenging |
| Iron storage/homeostasis | Iron | Ferritin-like storage and detoxification proteins |
| Zinc acquisition | Zinc | High-affinity Znu/Adc uptake and accessory zinc capture functions |
| Manganese acquisition | Manganese | MntH, MntABC-like, and SitABCD-like import systems |
| Copper homeostasis/resistance | Copper | P-type copper efflux, cue/cus branches, and copper chaperone/sensor systems |
| Cobalamin biosynthesis | Cobalt / vitamin B<sub>12</sub> | Anaerobic corrin-ring synthesis, late completion genes, and aerobic-route signatures |
| Cobalamin transport / cobalt uptake | Cobalt / vitamin B<sub>12</sub> | Btu transport, cobalt ABC uptake, and corrinoid salvage |
| Nickel utilization | Nickel | Nik/Hox/Nix transport with accessory maturation functions |
| Molybdenum cofactor biosynthesis | Molybdenum | Canonical moa/moe/mob pathway signals |
| Selenium utilization | Selenium | Selenocysteine and selenouridine utilization modules |

### 3.3 Genome-level demonstration on representative human-gut genomes

The MGnify demonstration provided a realistic genome-annotation use case. Across 11 representative human-gut genomes, copper homeostasis/resistance (mean score 0.861) and iron acquisition (0.839) were the most prevalent high-scoring programmes. Zinc acquisition (0.517), cobalamin transport/cobalt uptake (0.485), and cobalamin biosynthesis (0.444) followed at lower mean levels. Iron acquisition and copper homeostasis/resistance exceeded the calling threshold in all genomes in this panel, whereas zinc acquisition was called in 54.5% of genomes. The most variable phenotypes were iron storage/homeostasis, zinc acquisition, manganese acquisition, and nickel utilization, consistent with uneven lineage-level distribution of these programs.

Coarse phylum-group summaries showed biologically interpretable contrasts. Firmicutes displayed the strongest mean cobalamin-biosynthesis signal (0.613) together with clear selenium-utilization (0.427) and molybdenum-cofactor (0.377) signals. The Bacteroidota subset showed higher zinc acquisition (0.639) and manganese acquisition (0.400) but no detectable molybdenum cofactor or selenium signal in this small panel. The mixed *Other* group had the strongest cobalamin transport/cobalt uptake signal (0.667). No group comparison reached a low false-discovery rate threshold, which is unsurprising for only 11 genomes, but the smallest exploratory q values were observed for selenium utilization, cobalamin transport/cobalt uptake, molybdenum cofactor biosynthesis, and cobalamin biosynthesis.

Figure 1 shows the manuscript-style overview assembled directly from the TracePheno publication bundle for the MGnify run.

**Figure 1:**
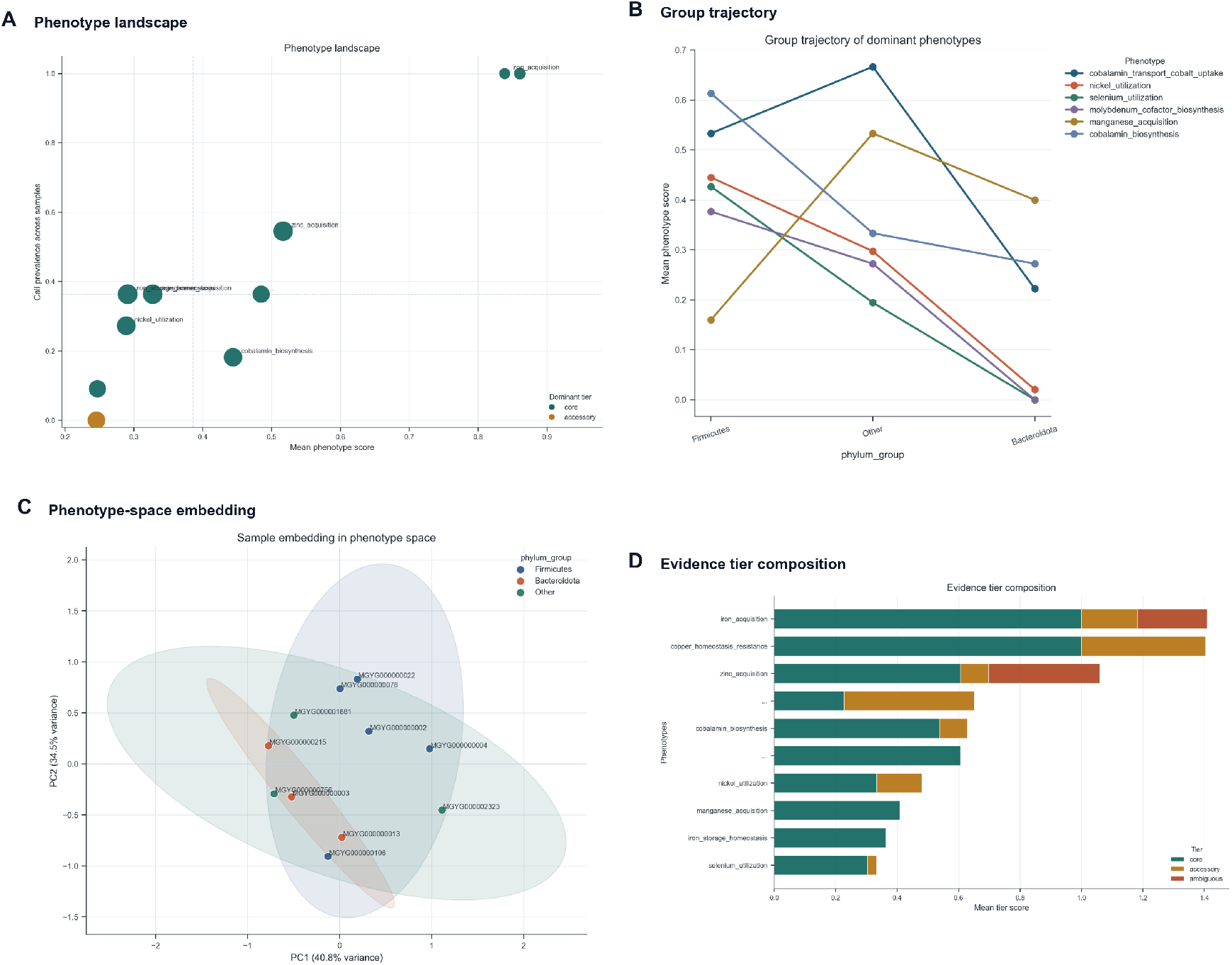
Global trace-element phenotype landscape. Overview panels curated for manuscript-style presentation from TracePheno outputs. Manuscript-style overview generated from the MGnify representative-genome demonstration (*n* = 11 genomes). Panel A summarizes the phenotype landscape by mean score, prevalence, variability, and dominant evidence tier; panel B highlights shifts across coarse phylum groups; panel C projects genomes into phenotype space; and panel D summarizes the balance of core, accessory, and ambiguous evidence.

### 3.4 PICRUSt2 compatibility enables a 16S-derived entry point

Although TracePheno is designed to avoid taxon-only phenotype transfer, many microbiome studies still begin with 16S data. The dedicated PICRUSt2 interface therefore provides a practical compromise: functional hypotheses are first projected to KO space by PICRUSt2 [Douglas et al., 2020], and TracePheno then maps those KO abundances to trace-element phenotypes. In the four-sample proof-of-concept example, the highest mean predicted phenotypes were zinc acquisition (0.567), cobalamin transport/cobalt uptake (0.535), iron acquisition (0.528), and selenium utilization (0.352). Binary calls were most prevalent for iron acquisition and zinc acquisition, each positive in three of four samples.

In the illustrative case-control split, zinc acquisition was higher in the case group (0.792 versus 0.341), whereas iron acquisition (0.723 versus 0.334), cobalamin transport/cobalt uptake (0.735 versus 0.334), and selenium utilization (0.477 versus 0.228) were relatively higher in controls. With only two samples per group, no phenotype survived multiple-testing correction, and the associated effect-size plots should therefore be interpreted as exploratory. Nonetheless, the example demonstrates that TracePheno can convert KO prediction tables into a biologically themed phenotype summary without requiring users to redesign the input matrix by hand.

### 3.5 Publication-oriented reporting extends beyond standard heat maps

Many microbiome software papers stop at matrix scoring and a single clustered heat map. TracePheno instead emits a coordinated set of figures designed for manuscript assembly. These include a phenotype landscape atlas, a differential phenotype map, raincloud-style group panels, clustered heat maps, and marker-contribution summaries, which are then assembled into composite figures with synchronized legends and summary text. The same run also produces a results-summary file that can serve as a scaffold for a narrative Results section.

Figure 3 illustrates the differential-statistics bundle for the PICRUSt2 demonstration, where the effect-size plot, group-mean heat map, and raincloud panels are arranged as a submission-ready composite.

**Figure 2:**
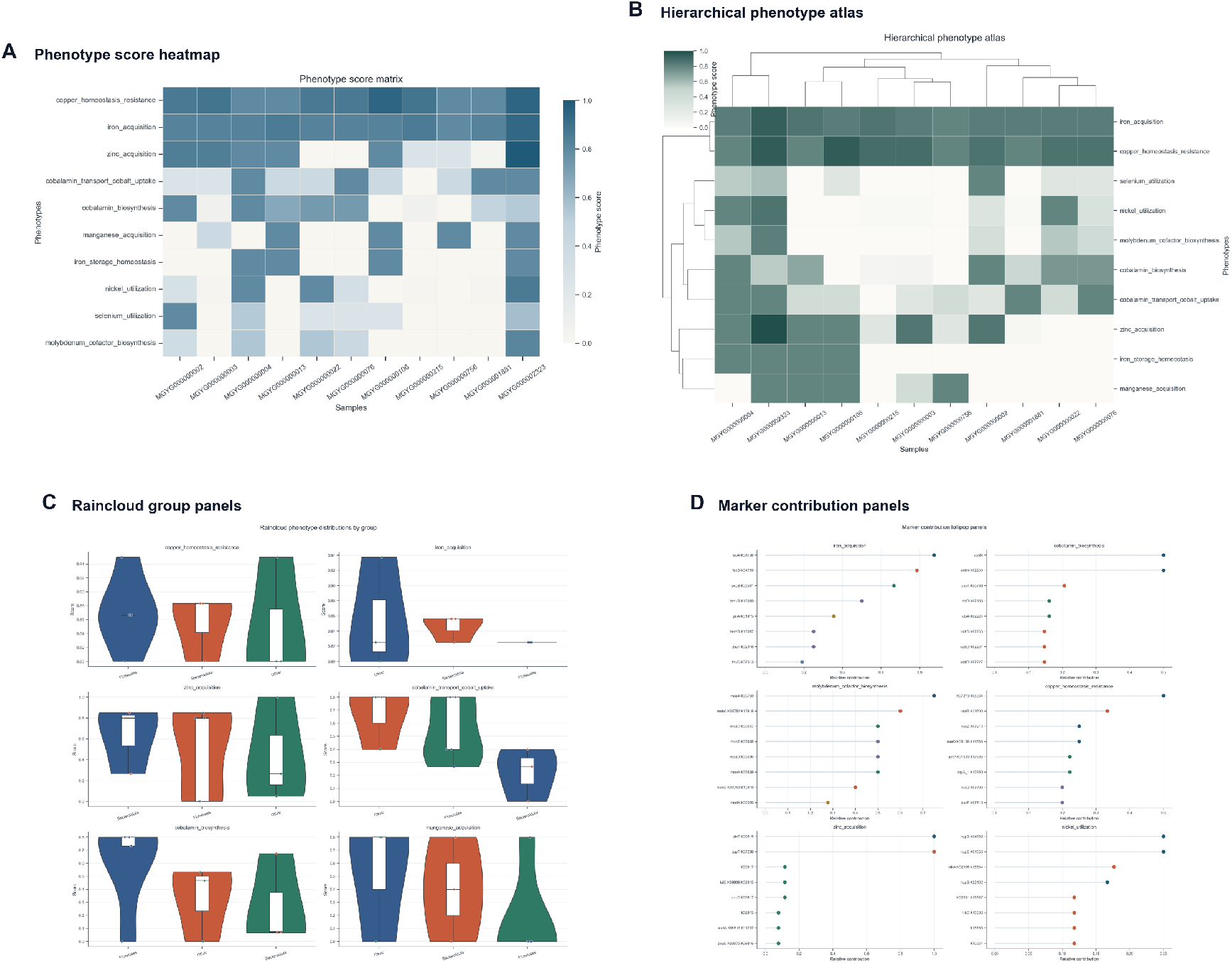
Cohort structure and phenotype drivers. Detailed panels that connect sample-level patterns to the markers supporting each phenotype. Structure-and-driver summary for the MGnify representative-genome demonstration. The multipanel figure links sample-level phenotype structure to the heat-map view, clustered phenotype atlas, groupresolved distribution patterns, and the marker contributions that support dominant phenotype calls.

**Figure 3:**
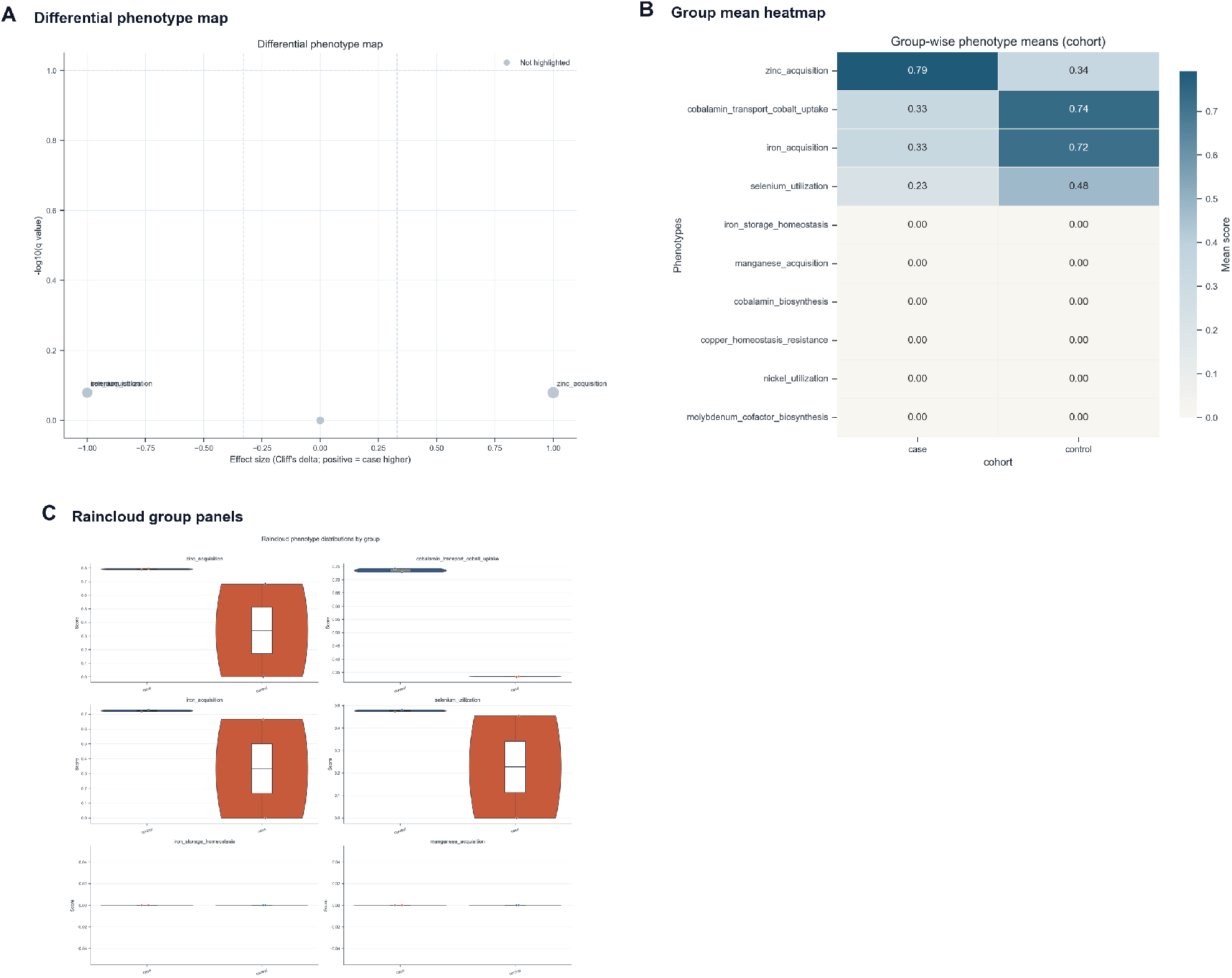
Differential phenotype statistics. Panels emphasizing cohort differences and effect-size-driven interpretation. Differential-statistics bundle generated from the PICRUSt2-style demonstration (*n* = 4 samples). The panel combines an effect-size map, group-wise mean heat map, and raincloud distribution panels. In this toy example, the figure is intended to demonstrate the workflow and visual grammar of the output rather than to support definitive biological inference.

## 4 Discussion

TracePheno was developed to occupy a methodological gap between taxonomy-oriented phenotype transfer and generic functional enrichment analysis. Its main conceptual contribution is to treat trace-element metabolism as a phenotype space that should be inferred from curated functions, not assumed from taxonomy alone. The current implementation sharpens that idea by making the decision rules explicit: marker evidence is bounded within samples, binary calls require predefined core evidence, multi-block pathways can demand multiple core modules, and genome trait inference can retain uncertainty rather than forcing missing evidence to zero.

The present demonstrations also clarify what the tool is and is not intended to do. In the genome-based example, TracePheno exposed broad copper and iron programs together with lineage-structured variation in cobalamin, molybdenum, nickel, and selenium functions. In the PICRUSt2 example, it provided a compact phenotype interpretation layer for KO predictions that would otherwise remain difficult to summarize biologically. At the same time, the software does not remove the uncertainty inherent to predicted functions. Any PICRUSt2-based analysis still inherits the limitations of 16S-to-function projection [Douglas et al., 2020], and the current implementation does not yet propagate NSTI-like uncertainty directly into phenotype confidence.

Several limitations remain. First, the phenotype library is curated rather than exhaustive, and extensions for arsenic, mercury, cadmium, chromium, and other metal systems still need to be benchmarked carefully before release. Second, although the current scoring rules are more interpretable than cohort-dependent threshold scans, they still rely on curated marker definitions rather than on direct experimental phenotype labels. Third, while quality-aware handling of incomplete genomes is available conceptually and algorithmically, broader benchmarking against experimentally characterized isolates and curated MAG collections will be needed to calibrate confidence more rigorously. Future work should therefore focus on library expansion, NSTI-aware uncertainty propagation for PICRUSt2 mode, and benchmarking against larger genome and metagenome collections with independent phenotype evidence. Even in its current form, however, TracePheno provides a more explicit bridge between microbiome function tables and element-resolved phenotype interpretation than either taxonomy-only heuristics or generic pathway summaries alone.

## 5 Software Availability

The current TracePheno implementation is publicly available through the [TracePheno GitHub repository](https://github.com/zjdevo/TracePheno). The phenotype definition files, demonstration datasets, and publication-style figures used in this manuscript are distributed with the software workspace and the associated repository contents referenced here. This manuscript draft directly cites figures generated by the local TracePheno runs.

## Acknowledgments

This manuscript draft was prepared from the current TracePheno implementation and its bundled demonstrations. This work was partially supported by the Postdoctoral Fellowship Program of CPSF (No. GZB20240728) and the China Postdoctoral Science Foundation Funded Project (No. 2024M763220).

